# Macroscopic and microscopic study on floral biology and pollination of *Cinnamomum verum* Blume (Sri Lankan)

**DOI:** 10.1101/2022.07.12.499711

**Authors:** Bhagya Madhushani Hathurusinghe, D.K.N. Gamini Pushpakumara, Pradeepa C.G. Bandaranayake

**Author notes:** Correspondence to P.C.G. Bandaranayake.

## Abstract

*Cinnamomum verum* Blume (syn *Cinnamomum zeylanicum*) commonly known as Ceylon cinnamon, has gained worldwide attention and well supported by experimental data. Maintaining the yield quality and quantity are essential, especially for high-end value-added products. Breeding superior varieties and producing good quality planting materials are critical for commercial cultivation efforts. However, limited literature is available on the floral development and biology of *C. verum*. We assessed the seasonal flowering, floral development and pollination of type A and type B varieties of *C. verum*. Both macroscopic and microscopic data were collected on floral biology, pollination, male and female floral organs before and after pollination. *Cinnamomum verum* is morpho-anatomically, structurally, and physiologically adapted for cross-pollination, possible between the two cultivars; type A (*Sri Gemunu*) and type B (*Sri Wijaya*) flowers; naturally evolved with Protogynous Dichogamy. It possesses a dry stigma, which changes its surface characteristics after pollination. During close-pollination, a higher percentage of pollen shrink and shrivel on stigma while it develops callose plugs to obstruct the growth of the pollen tube. However, during changes in environmental conditions, female and male stages in the same tree overlap for about 45-60 min suggesting possible close-pollination within the same plant. Moreover, 4-8% of the flowers formed fruits after natural close and hand pollination. pollen was found fully hydrated even after close-pollination. Although *C. verum* is adapted for cross-pollination, natural close-pollination is also possible. The data suggest the complex nature of the sexual reproduction of *C. verum*. Well-managed breeding attempts will develop superior *C. verum* varieties.

## INTRODUCTION

The Genus *Cinnamomum* Scharffers belongs to the family Lauraceae, which comprises over 250 species [1] distributed from the Asiatic mainland to Formosa, the Pacific islands, Australia, and Tropical America [2]. *Cinnamomum* is a pantropical genus consisting of evergreen trees and shrubs [3]. While some economically important species such as *C. camphora* J. Prei and *C. aromaticum* Nees are trees, others such as *C. verum, C cassia, Cinnamomum burmannii*, (Indonesian cinnamon), and *Cinnamomum loureiroi* (Vietnamese cinnamon) [4], and *Cinnamomum tamala* T.Nees & C.H.Eberm., (India and Nepal) [5] are shrubs.

All the *Cinnamomum* species recorded so far are cross-pollinated with Synchronised Protogynous Dichogmay as the prominent mechanism [6,7]. Dicogamy is defined as the temporal separation of male and female functions. In protandrous species, another dehiscence occurs before the stigma becomes receptive, while it is reversed in protogynous species [8]. In some species, the same flower opens twice, either in the morning or in the afternoon of the first day, and the afternoon or the morning of the next day, respectively [9]. The same phenomenon has been recorded in *C. verum* in Sri Lanka [10].

Avocado, one of the other economically important species in the family Lauraceae, also evolved with Protogynous Dichogamy breeding system [11]. The Synchronous dichogamy of avocado presents two morphs for successful cross-pollination and flowering is complementarily synchronized. However, the flowering is extremely sensitive to environmental conditions [12,13], where flowering behaviour fluctuates with temperatures and humidity [11,14]. Davenport in 1986 defined the morphological development of the avocado flower, relating to the histological details. However, such information on *Cinnamomum* is limited.

Azad and colleagues discussed the morphological variation of the flowers and inflorescences and mutations in *C. verum* [16]. The cinnamon flower has both functional male and female organs [17], and there are two types of varieties. In the type-A varieties, female flowers open for the first time in early or mid-morning, remain open and pistil receptive until about noon, then close and remain closed until noon of the following day, when they reopen and begin shedding pollen with the pistil no longer receptive. The flowers close permanently in the late evening, after a cycle of about 36 hours. The flowers of type B varieties function analogously but with transposed timing. The opening cycle of type B flowers spans about 24 hours, and the difference in cycle time reflects the relative length of the closed period between openings [17]. Therefore, cross-pollination is possible with type A and type B varieties. Since vegetatively propagated planting materials create crop management issues in largescale cultivations [18,19], seedlings are preferred.

For hybridization efforts, optimization of cross-pollination and avoiding possible close-pollination are vital. On the other hand, maximizing close-pollination would benefit the planting material production from identified superior genotypes. A closer look at floral biology, pollination, and fertilization would facilitate such efforts. Therefore, the current study focused on the detailed floral biology and morphology of selected type A and type B of *C. verum* varieties before and after pollination.

## MATERIALS AND METHODS

### Study site and sample collection

The experiments were conducted for two consecutive years (2019-2020) in a Vegetative propagated orchard located at the Delpitiya sub research station of the Department of Export Agriculture, Atabage, Sri Lanka (707.970’N, 80035.342’E; 634m MSL) and greenhouse at the Agricultural Biotechnology Centre, Peradeniya, Sri Lanka. The experimental field consisted of two verities *Sri Gemunu* (Type A) and *Sri Wijaya* (Type B), in alternative rows. Six plants with flowers from each variety were marked and selected as replicates. The flowering phenology and floral behaviour were monitored from 7.30 am to 4.00 pm. The meteorological data viz., minimum and maximum temperature and relative humidity of Delpitiya were recorded. The temperature was recorded hourly while the humidity was recorded two times a day.

### Flower and inflorescence morphology

Key morphological attributes of flowers and inflorescences were measured to characterize the visual display and the resource accessibility of the reproductive units. The length of flowering inflorescences was determined in the field on selected individuals with a ruler. The number of flowers in inflorescences, and the number of inflorescence in inflorescence clusters were counted in the field. Three selected inflorescences clusters were tagged. We recorded the number of opened and closed flowers in an inflorescence, flowering period, and flower lifespan, and measured the length of the stigma, petal, stamen, and corolla taking the mean values of 10 replicates. Floral morphology was studied with fresh flowers using a Stereomicroscope (Kruss Opironic and Olymus SZX10). The images were captured from the software AmScope V/ 3.7.2776 and Oylmus Cell Sens Standard V 1.16). Dimensions of single floral organs and their position within the flower were determined with size differenced micrographs using the ImageJ software [20].

### Dynamics of Flowering

Six trees from both cultivars, *Sri Gemunu* and *Sri Wijaya*, were marked. In 2019 and 2020, we collected flower phenology data by counting the number of flowers open in each selected plant during both phases. The length of the flowering period was defined as the number of days between the opening of the first flower and the closing of the last flowers. It was measured for two years in 2019 and 2020, during the mass flowering season, January-February. Flowers were considered open if pollinators could use their petals at least 45 degrees open and closed if both the stigmas and anthers wilted. The flowering progress was monitored at 30-minute intervals every day. We used the phenology data collected to determine the mass or peak flowering season for each variety, flowering period, and overlapping period. The flowering of labelled inflorescence was observed for 18 days during flowering seasons before flowering, with 25% of flowers open, peak flowering >50% open and termination of flowering. To determine the flowering periods, the number of pollination units with opened flowers in each plant was labelled and counted.

Flowering synchrony was measured using an index that considers the proportion of open female flowers on the focal plant that overlapped temporally with the open male flowers in the next phase. The synchrony index (SI) was calculated with the equation given [21];

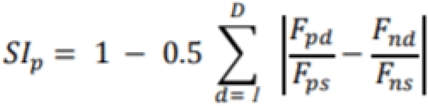

The index outputs values from zero to one, where one is the most synchronous and indicial could be and zero is not synchronous at all. Synchrony index was counted for the flowers that were open during the overlap period.

where *F*_*pd*_ and *F*_*nd*_ are the numbers of flowers open on focal plant *p* and all of its neighbours *n*, respectively, on day *d*; *D* is the number of days those flowers were counted for an individual; and *F*_*ps*_ and *F*_*ns*_ are the total numbers of flowers produced during the season by focal plant *p* and all of its neighbours *n*, respectively.

### Pollen-Stigma Interaction

We conducted controlled experimental pollinations to understand the possibility of natural selfing in *C. verum*. Three plants of *Sri Gemunu* were isolated in separate net houses made from shade nets/mesh (Hole size 1 mm x 2 mm). They were built to avoid the possibility of pollination from the neighbour plant and to avoid any external influence on pollination and kept at a considerable distance (Fig 1). Out of the three plants, two were allowed to close-pollinate via ants, which was identified as an efficient pollinator in the field, while another one was close-pollinated by hand. Further, five (5) pistils from each pollinated flower at different time intervals (after 1 hour, 4 hours, and 20 hours of opening) were collected to observe the growth of the pollen tube. Pistils were immediately subjected to fixing solution (acetic acid: absolute alcohol, 1:3) overnight and then transferred to70% ethanol and preserved.

**Fig. 1:**
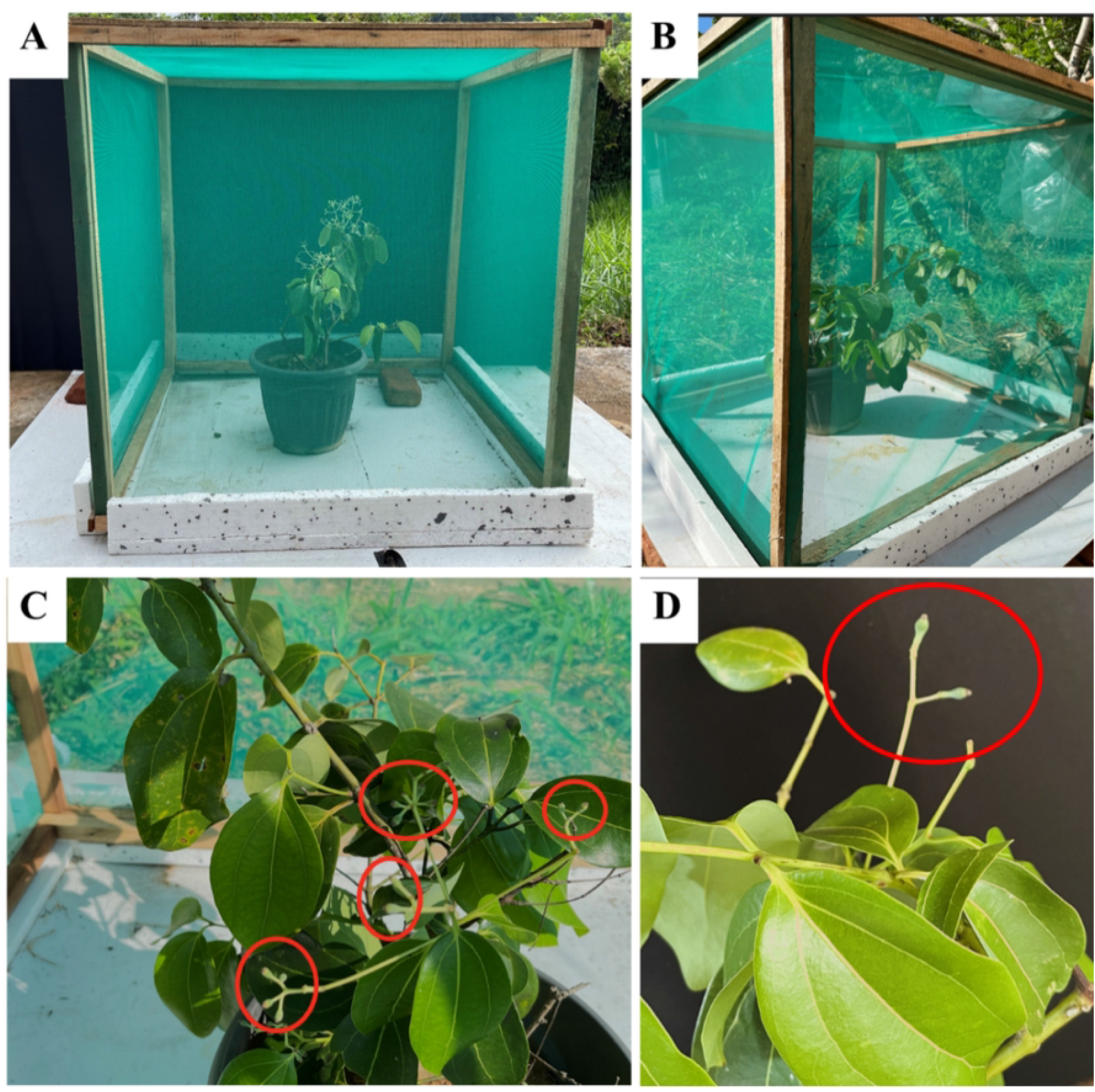
Field experimental blocks for observing the efficiency in close pollination located at Agricultural Biotechnology Centre. (A) Net house built for isolation of cinnamon plants for insect pollination and hand pollination. Insect pollination is from ants (B) Net house built for isolation of cinnamon plants for hand pollination. hand pollination was done between two female and male flowers from the same tree, during the overlapping period (C) fruit production yielding 8% of total flowers produced, pollination method: hand pollination (D) fruit production yielding 4% of total flowers produced, pollination methods: insect pollination

To observe the pollen and stigma interaction, the flowers were harvested from three plants at three stages, i. female flowers just opened before being stigma receptive, ii. female flowers after stigma being receptive iii. female flowers after being pollinated, and iv. pollinated male flowers after anthesis. Three individual flowers from each were selected, and visibly observed pollination by the pollinators after 3-5 hours of observation.

The structural and morphological variation in the stigma was observed before and after pollination in both ‘*Sri Wijaya*’ and ‘*Sri Gemunu*’, using a Scanning Electron Microscope. Further stigma-pollen interaction and self (in)compatibility were observed using a fluorescence microscope.

### Scanning Electron Microscopy, sample preparation and Imaging

The pollen and stigma of the collected specimens were studied using Scanning Electron Microscopy (SEM). For SEM analysis, the flowers at the anthesis stage and pollinated stigma were used. The fresh anthers and stigma were separated from the flower and fixed in FAA solution, 5% formaldehyde (v/v), 5% (v/v) acetic acid, and 45% (v/v) ethanol for long term preservation.

The preserved fresh pollen samples were centrifuged at 2,500 rpm for 3 minutes. The pellet was then washed twice with distilled water for 25-30 minutes. After the washing, pollens were dehydrated in an ethanol series (100% DW→3:1→1:1→3:1→100% ethanol) for about 15 minutes at each stage. The pollens were separated by centrifugation at 2500 rpm for 3 minutes at each step, after making sure that the ethanol and water are removed. For specimen drying, the samples were then transferred into 100% Hexamethyldisilazane (HMDS) through a graded series (100% ethanol→3:1→1:1→3:1→100% HMDS) and soaked for about 20 minutes at each stage. Then pollen was air-dried until all HMDS get evaporated. The processing of the preserved stigmas followed the same chemical treatments without centrifugation steps.

The samples were mounted onto the sample stub using carbon tapes and gold sputter-coated for 15 seconds. The coated samples were examined by Hitachi SU6600 Analytical Variable Pressure FE-SEM (Scanning Electron Microscope) at Sri Lanka Institute 0f Nanotechnology (SLINTEC), Homagama, Sri Lanka.

### Fluorescence Microscopy

We followed the protocol described by Dafni in 1992 [22]. For the observation of pollen tubes growing inside the style, the fixed excised stigma and style in FAA were treated with 70% and then were washed with distilled water. Fixed stigma samples were kept at +4 2°C in the refrigerator until staining. Pistils were softened by keeping them in 8 M NaOH at 60°C for 4h. They were then rinsed in running water for 3 hours to remove NaOH. Then stained in 0.005% aniline blue in potassium acetate for 6 h and mounted in a 1:1 v/v mixture of aniline blue and glycerin. The slides were observed under a fluorescence microscope with a UV filter and the germination of pollen on the stigma and path of pollen tubes through the stylar tissue to the ovule.

### Statistical analysis

We checked the data distributions for normality using Sharpiro-Wilk normality test before selecting parametric or nonparametric variations of each statistical test. Variances of data distribution were compared using F-tests. For analyses comparing the two groups *Sri Gemunu* and *Sri Wijaya*, we used Mann-Ehitney U tests, whereas t-tests were used for groups with equal variances and normal distributions. One-way analyses of variance were used to test differences among the populations, and their significance was tested through Fisher’s least significance difference (LSD). All analyses were done in SAS studio version 3.8 and Analyse-it Standard Version 2.30.

## RESULTS

### Floral Biology

*C. verum* flowers are actinomorphic with long greenish-white colour peduncles (Fig 2 A), bracteate, bisexual, trimerous (Fig 2 A, B), and perigynous. In the front aspect, single flowers of each (or both) sex are seen as clear to yellowish-green. Tepals on the outside are greenish-white oblong-lanceolate, tepals on the interior are pale white (Fig 2 A, Fig 3), and newly opened flower nectaries are yellow (Fig 2 A,B). There were no distinct floral morphology differences between *Sri Gemunu* and *Sri Wijaya*.

**Fig. 2:**
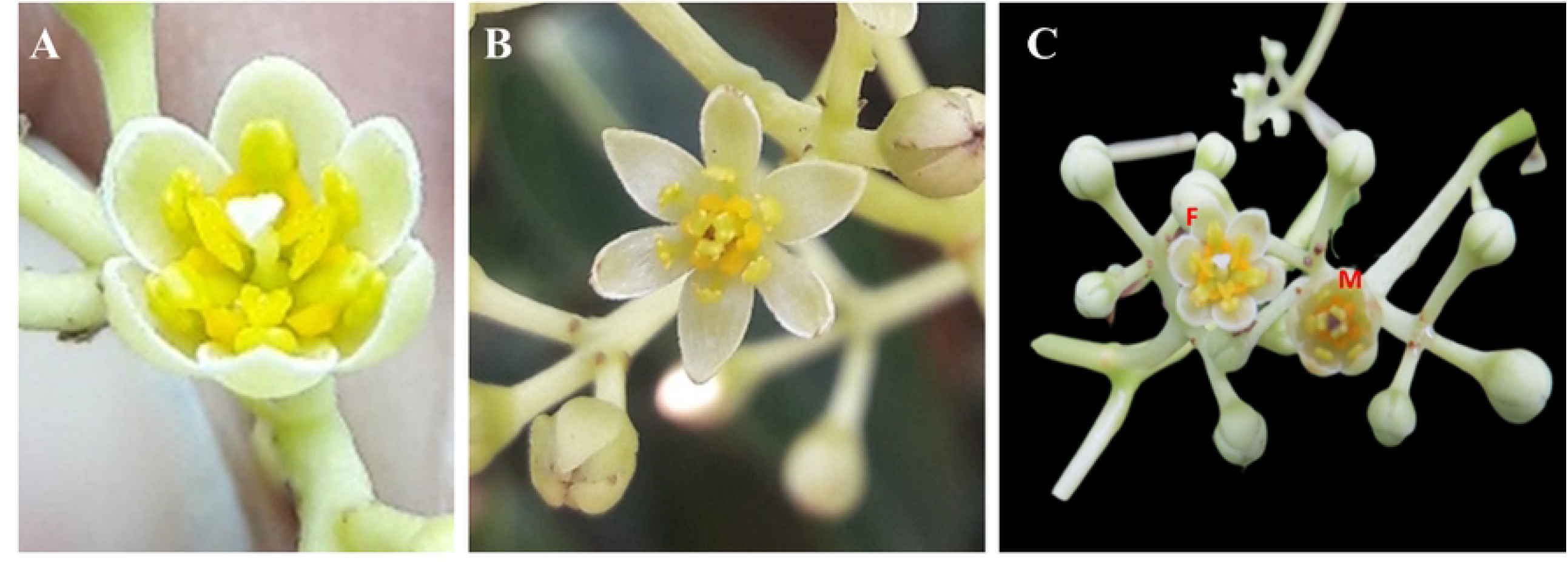
Floral Morphology of Female and Male flowers. (A) fully opened female flower (B) fully opened male flower (C) female and the male stages opened in the same tree which can result in close-pollination

**Fig. 3:**
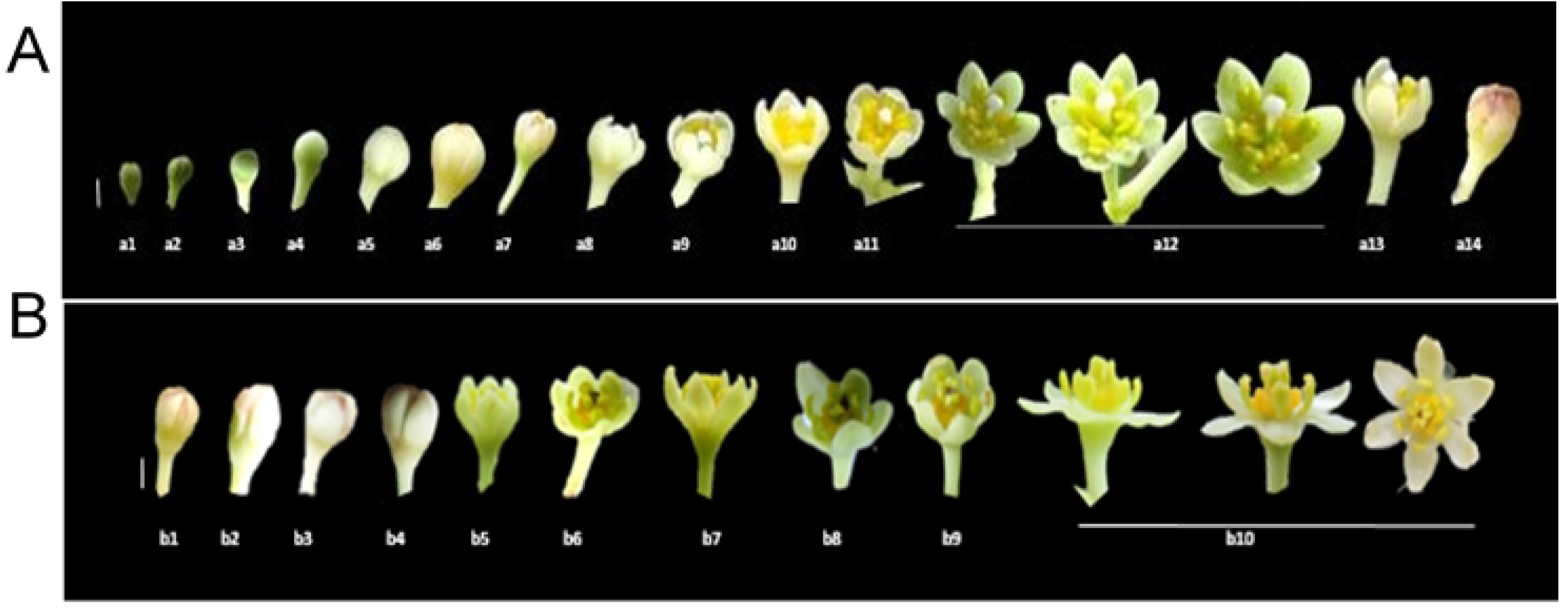
The sequential floral development of cinnamon; illustrating the development of newly emerged flower bud to fully developed female flower and subsequently to male flower. (A) a1-a14 Changes of flowering in the female flower during the first phase. a1-a4 immature flower bud with light green surface; a5 small-sized bud with light green surface; a6-medium-sized mature bud with yellow surface; a7-a9-Initiating bloom; a10-a11-half blooming stage; a12-fully blooming stage; a13-a14 wither stage (a bar-1 mm) (B) b1-b10 Changes of flowering in the male flower during the second phase. b1-b4 initial flower bud with pale colour; b5-b7-initiating bloom; b8-b9-half blooming stage; b10-fully blooming stage (b bar – 2 mm)

### Flower development of type A and Type B cultivars

n *Sri Gemunu* cultivar, flowers open for the first time in early or mid-morning, remain open and the pistil is receptive until about noon. Then the flower closes and remains closed until about noon on the second day. When those reopen, begin shedding pollen with the pistil no longer receptive. Finally, those close permanently on the second day evening. The complete opening and closing cycle of *Sri Gemunu* spans about 36 hours. On the same tree, many flowers open for the first time in the same morning and then follow the same behaviour pattern synchronously hour after hour for their 2-day existence. The flowers of the *Sri Wijaya* cultivar function analogously but with transposed timing. The opening cycle of *Sri Gemunu* spans about 24 hours. The difference in cycle time reflects the relative length of the closed period between openings. Anther dehiscence occurs after 30 minutes to 1 hour from the second flower opening. The surface of the stigma turns brown and shrivelled by the time pollen is released (Fig 3).

#### Female stage

The female stage flower is half open with a white stigma that stands erect and separate, ready to receive pollen (Fig 4 A, C, D) though the pollen is not released from the closed pollen sacs of that flower (Fig 4 F). The stigma was tiny, papillose, and slightly capitate; the single uni-ovulate carpel was associated for up to 80% of the ovary and plicate above this and along with the elongate style (Fig 4 G, H). Female cinnamon flowers have receptive stigmata, silky white (Colour code: 1 A 1; colour values according to Kornerup and Wanscher 1981) (Fig 4 H). The gynoecium is produced by a unilocular pistil and contains one ovule (Figure 5 H). Stamens are free, 9+3 in four whorls in three each (Fig 4 A, B). The anthers are with unopened pollen sacs at the end of the stamens (Fig 4 J). It has all 12 stamens bent at almost a 90-degree angle to the central erect pistil (Fig 4 D, 1 A). Nectar is secreted from the three staminodes located at the base of the third whorl of stamens (Fig 4 E, F, I).

**Fig. 4:**
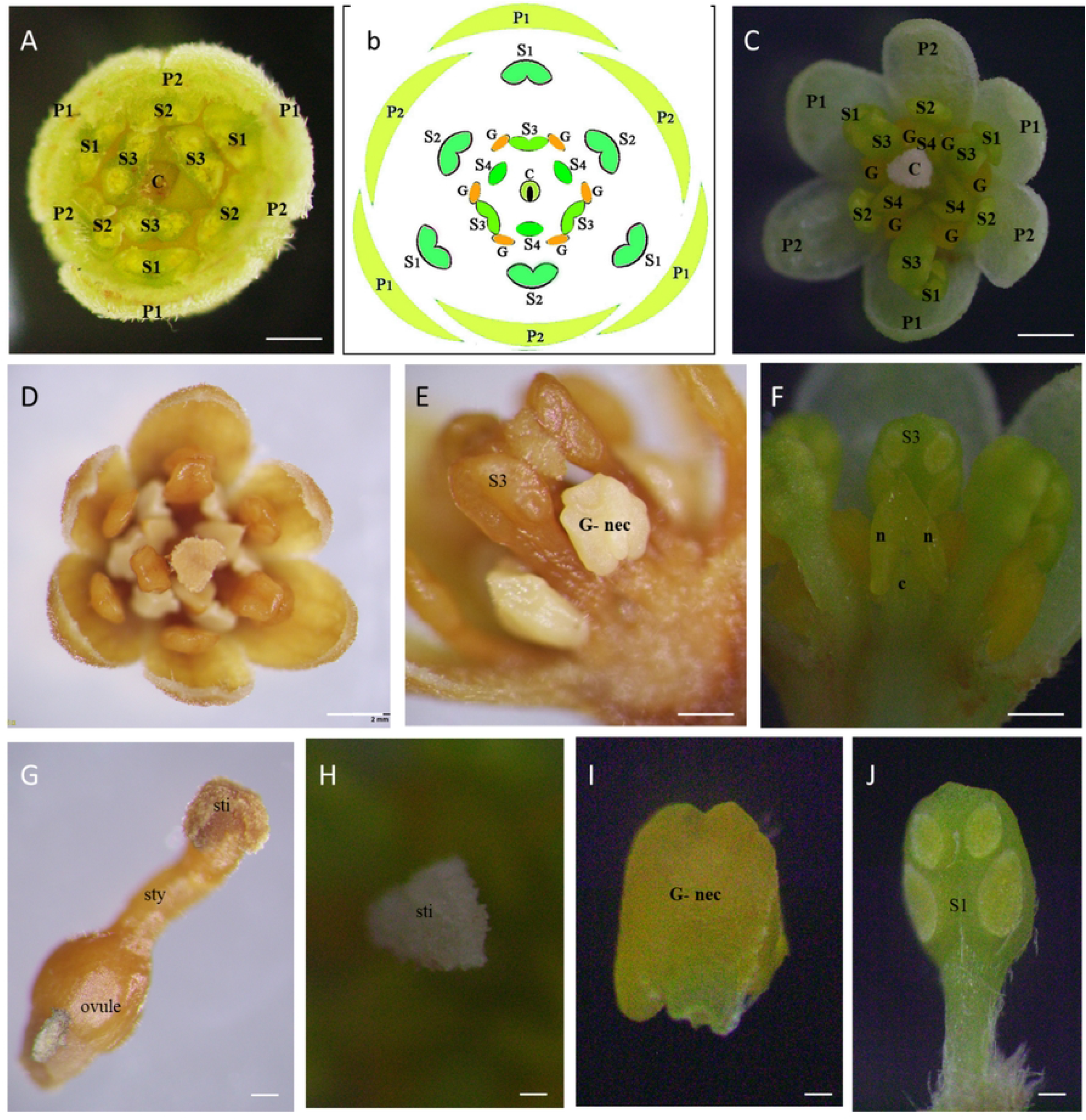
Flower and floral organs of *Cinnamomum verum* functional female flower. (A) Transverse section of flowers (B) Schematic representation of assumed floral organization in *Cinnamomum verum* flower (C,D) Flower at the female stage, apical view, dissecting microscopy (E) Stamen of the third whorl during the female stage, with a pair of paired staminal appendages, abaxial view (F) Slightly aberrant staminode (fourth androecial whorl) before anthesis, with lateral nectary tissue separated by a constriction, reminiscent of pollen sacs, apparently separated by a prominent mid-portion connective, adaxial view (G) Pistil, ovary in longitudinal-section, showing single, apical ovule (H) Dorsal ventral view of female receptive stigma, milk-white triangular shaped (I) adaxial view of nectariferous structure without anther (J) adaxial view of the unopened stamen (S1) Abbreviations. C, carpel; G, gland; P1, the outer whorl of tepals; P2, the inner whorl of tepals; S1, the first/outermost whorl of stamens; S2, the second whorl of stamens; S3, the third androecial whorl including either fertile stamens or staminodes; S4, the fourth androecial whorl including staminodes.

**Fig. 5:**
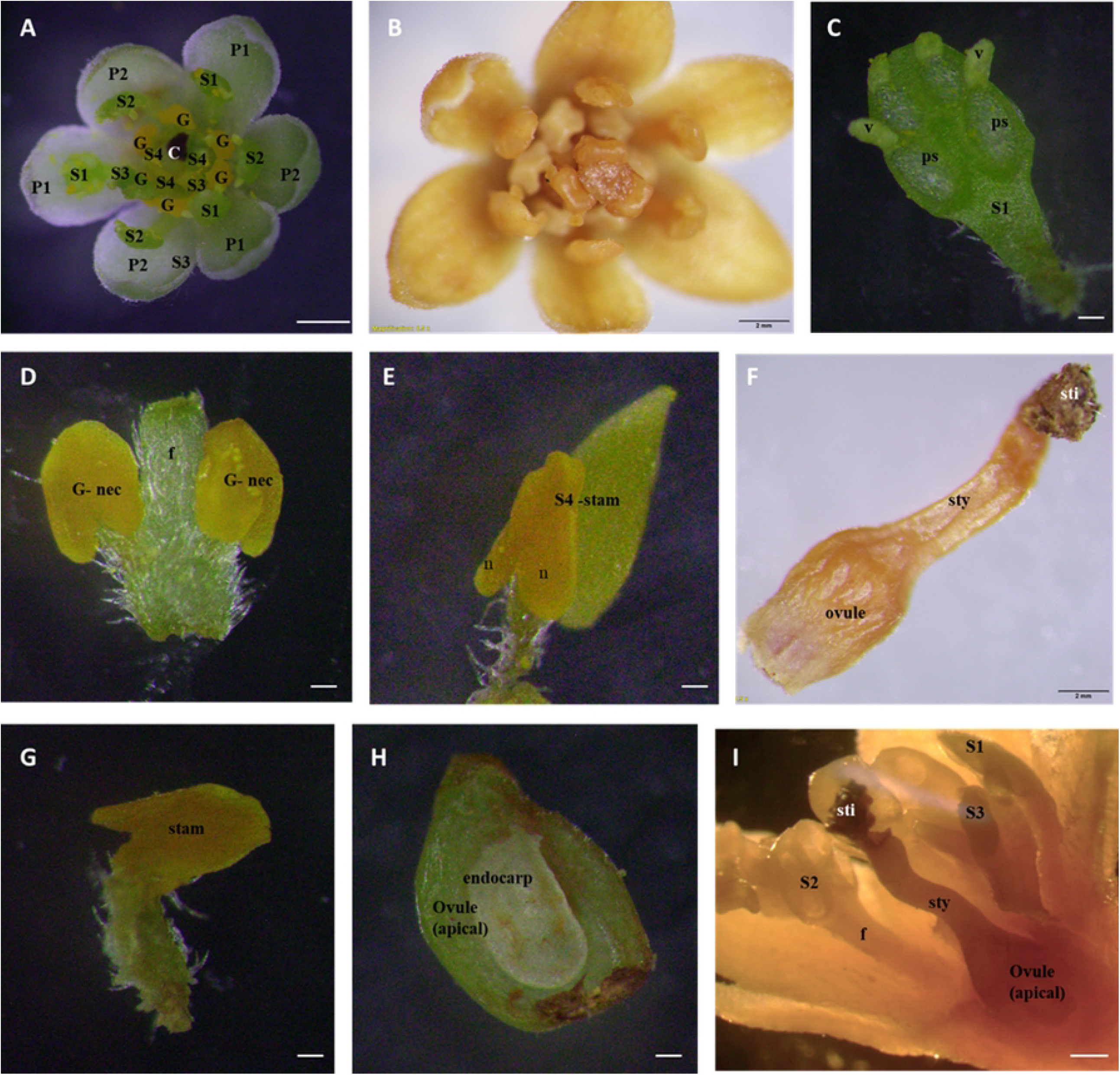
Flower and floral organs of *Cinnamomum verum* functional male flower. (A,B) Transverse section of male flower, at anthesis (C) First whorl stamen at anthers with open stoma flaps, the base has no trace of additional structures, adaxial view (D) Close up of pair of appendages at the base of a third whorl stamen, abaxial view (E) Aberrant inner tepal at anthesis, with yellow, putatively nectariferous structure at the base, with slightly aberrant staminode at anthesis, with lateral nectary tissue separated by a constriction (F) pistil, darkened stigma pollinated, with style and the ovary (G) the fourth whorl stamen lateral view, (H) ovary in longitudinal-section, showing single, apical ovule (I) lateral view of pollinated flowers with opened stamens in the three whorls and stigma darkened with fibrous-like structures

#### Male stage

The flower opens as a functional male after closing overnight after the female opening. The male flowers were fully open when the stigma turned brown or black and wilted at the center (Fig 2 B, Fig 5 A, B, I, F). The androecium has four trimerous whorls (floral whorls 3-6), with the outermost whorl alternating with the inner tepals; three outer whorls of fertile stamens and an inner sterile whorl of staminodes; and three outer whorls of fertile stamens, and an inner sterile whorl of staminodes (floral whorl 6, androecial whorl 4) (Fig 5 B). Mature stamens are longer upright, prominent, and sticky. Inner three stand erect in the middle around the pistil, and the other six stand at an angle of 500-600 closer to the pistil (Fig 5 A, B). Between the internal and exterior stamens, those form a ring at the base, identified as the floral nectary (Fig 5 D, E, G).

There is no obstruction to nectar entry other than the gap between internal and external stamens (Fig 5 D, E). The anther dehiscence occurs after 1-2 hours from the second opening of the flower. The surface of the stigma turns brown normally and shrivelled by the time pollen is released (Fig 5 I).

The stamens of the two outer whorls (androecial 1,2) were essentially identical and had a long, hairy filament as well as a cylindrical or box-shaped anther (Supplementary Fig 1). Each anther has four half-thecae with elliptical valves that only dehisced in male individuals’ flowers that opened introrsely. Only those individuals had pollen in their anther thecae, which was visible on the former inner side of the constructed valves (Fig 5 C). The extrorse anthers and the presence of a pair of basal staminal appendages distinguished the stamens of the following inner whorl (androecial whorl 3) from those of the outer whorls. Both appendages were placed at the filament’s base and had a white, hairy stalk-like filament (Fig 5 J).

Accordingly, the apices of the appendages are kidney-shaped, hairy, triangular, and comparable to the apical structure of staminodes at maturity. The appendage apex’s yellow bulge was more visible on the adaxial side (to the floral axis) (Fig 5 F, E). The staminodes (androecial whorl 4) had a shorter filament and a yellow swelling apex, triangular-acute with a tip akin to an apical connective appendage, and more noticeably bulging adaxially, with a larger connective on the abaxial side. Nectar was secreted by staminodes and paired staminal appendages.

The open *Cinnammomum* male and female flowers were compared. The male flowers were considerably larger (Table 1). The male stage flowers were fully opened, and the female stage flowers were concave-shaped. Significant differences were in the median flower diameter (male: 6.22 mm in SG 6.23 mm in SW vs. female: 6.09 mm in SG and 6.11 mm in SW), the mean stamen length (male: 2.4 mm vs. female: 2.1 mm), and the mean pistil height (male: 1.15 mm vs. female). Even though the female pistils and stigmata were higher than the male pistils, those were much taller than the surrounding anthers.

**Table 1:** Floral characteristics of *Cinnamomum verum* Sri Wijaya and Sri Gemunu.

| Floral characteristics |  | *Sri Gemunu* |  | *Sri Wijaya* |  |
| --- | --- | --- | --- | --- | --- |
|  |  | Female | Male | Female | Male |
| Flower length (mm) | Min-max | 6.8-10.3 | 7.3-11.1 | 6.75-10.6 | 7.35-11.45 |
|  | Ave±SD | 8.58±1.27 | 9.08±1.31 | 8.80±1.25 | 9.10±1.30 |
| Flower diameter (mm) | Min-max | 5.2-6.6 | 5.3-6.79 | 5.2-6.5 | 5.3-6.8 |
|  | Ave±SD | 6.09±0.39 | 6.22±0.40 | 6.11±0.37 | 6.23±0.39 |
| Pedicle length (mm) | Min-max | 0.73-1.2 | 0.73-1.2 | 0.75-1.5 | 0.75-1.5 |
|  | Ave±SD | 0.93±0.13 | 0.93±0.13 | 0.98±0.21 | 0.98±0.21 |
| Inflorescence length (mm) | Min-max | 3.5-17.4 | 3.5-17.4 | 3.8-17.4 | 3.8-17.4 |
|  | Ave±SD | 13.43±4.47 | 13.43±4.47 | 13.19±4.15 | 13.19±4.15 |
| Stigma diameter (mm) | Min-max | 0.25-0.62 | 0.24-0.59 | 0.26-0.58 | 0.28-0.60 |
|  | Ave±SD | 0.45±0.13 | 0.42±0.14 | 0.45±0.09 | 0.48±0.10 |
| Style length (mm) | Min-max | 0.91-1.8 | 0.78-1.5 | 0.9-1.75 | 0.80-1.49 |
|  | Ave±SD | 1.35±0.3 | 1.13±0.23 | 1.38±0.26 | 1.15±0.19 |
| Petal length (mm) | Min-max | 2.89-3.5 | 2.89-3.5 | 2.75-3.5 | 2.75-3.5 |
|  | Ave±SD | 3.13±0.18 | 3.13±0.18 | 3.14±0.21 | 3.14±0.21 |
| Petal width (mm) | Min-max | 1.2-1.64 | 1.2-1.64 | 1.35-1.63 | 1.35-1.63 |
|  | Ave±SD | 1.5±0.11 | 1.5±0.11 | 1.49±0.09 | 1.49±0.09 |
| First and second whorl anther length (mm) | Min-max | 1.8-2.23 | 2-2.55 | 1.92-2.25 | 2.03-2.6 |
|  | Ave±SD | 2.1±0.12 | 2.4±0.15 | 2.11±0.15 | 2.41±0.14 |
| Flowers per inflorescence | Min-Max | 28-158 |  | 18-216 |  |
|  |  | 81.5±45.61 |  | 101.52±70.38 |  |

### Flowering period, floral morph proportion and flower life span

The flowering of *C. verum* occurs mainly from the end of December to the middle of February. Flowering amplitude was the highest in the mass flowering period at the beginning of February (Fig 6 A and B). The total duration of the flowering period in a population was about one month. For an average flower, it takes 13-15 days for development from the day of initiation. The opening of the flowers occurred in two stages; the first opening was the female stage, and the second opening was the male stage. It looks receptive at the first stage after the flower opened, and there was no anther dehiscence (Fig 2 A). The flowers closed and opened again the next day, i.e., the second stage when the anther dehisces in the second time opening of the flower (Fig 2 B).

**Fig. 6:**
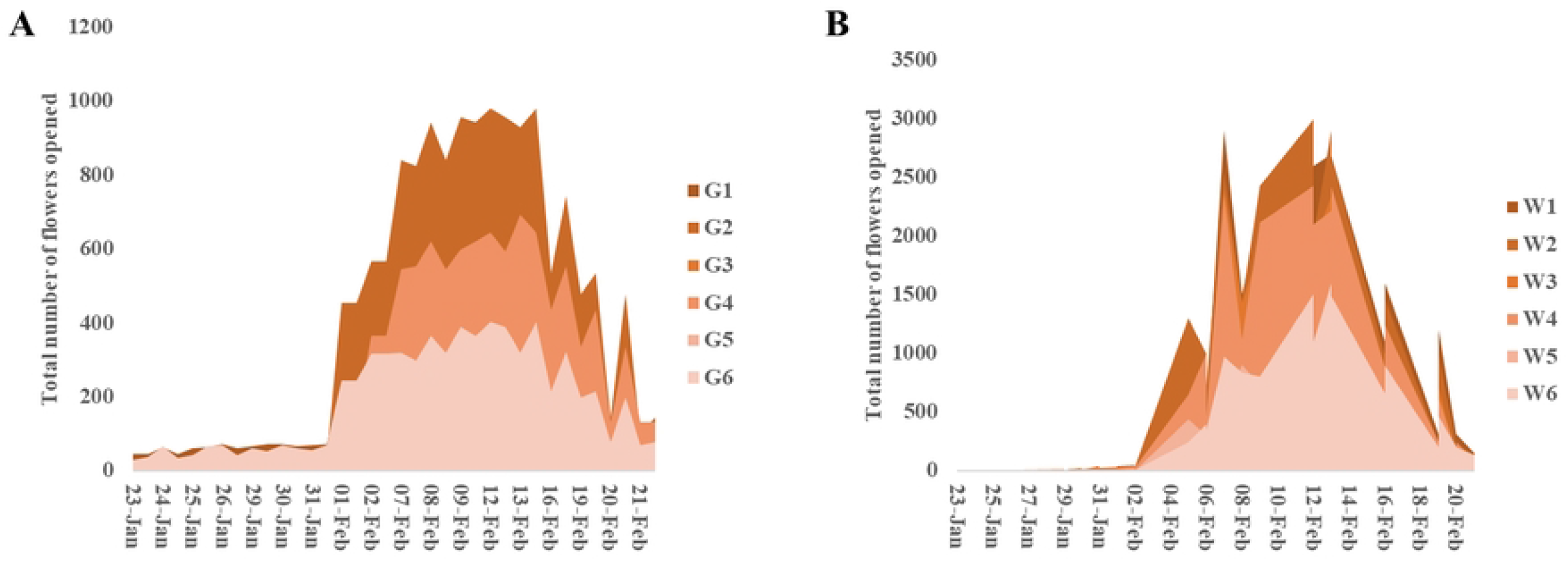
Progression of flowering in (A) *Sri Gemunu* and (B) *Sri Wijaya* at Deplitiya Sub research station in 2019.

### Floral Behavior: Overlapping Period

As a result of mass flowering, under ideal temperatures (maximum 32°C, minimum 27°C) (Supplementary Figs. 1, 2) there was some overlap with other flowers providing a small window of opportunity for close pollination. The maximum overlapping period was seen on the day having a temperature 29°C-32°C in both the varieties (Supplementary Figs. 2, 3). A long overlapping period of female and male flowers was observed in *Sri Gemunu* from around 10.00 am – 11.45 am, while in *Sri Wijaya* female and male flowers overlapped between 12.00 am-12.35 am (Fig 7 A). Based on our results, the optimum temperature and humidity for overlapping during mass flowering were 29°C-30°C and >70%, respectively (Supplementary Fig 2, 3) It provides a small window of opportunity for close pollination. We found an overlap of 45 min to 90 min in *Sri Gemunu*, while it was 45 minutes in *Sri Wijaya* (Fig 7 B).

**Fig. 7:**
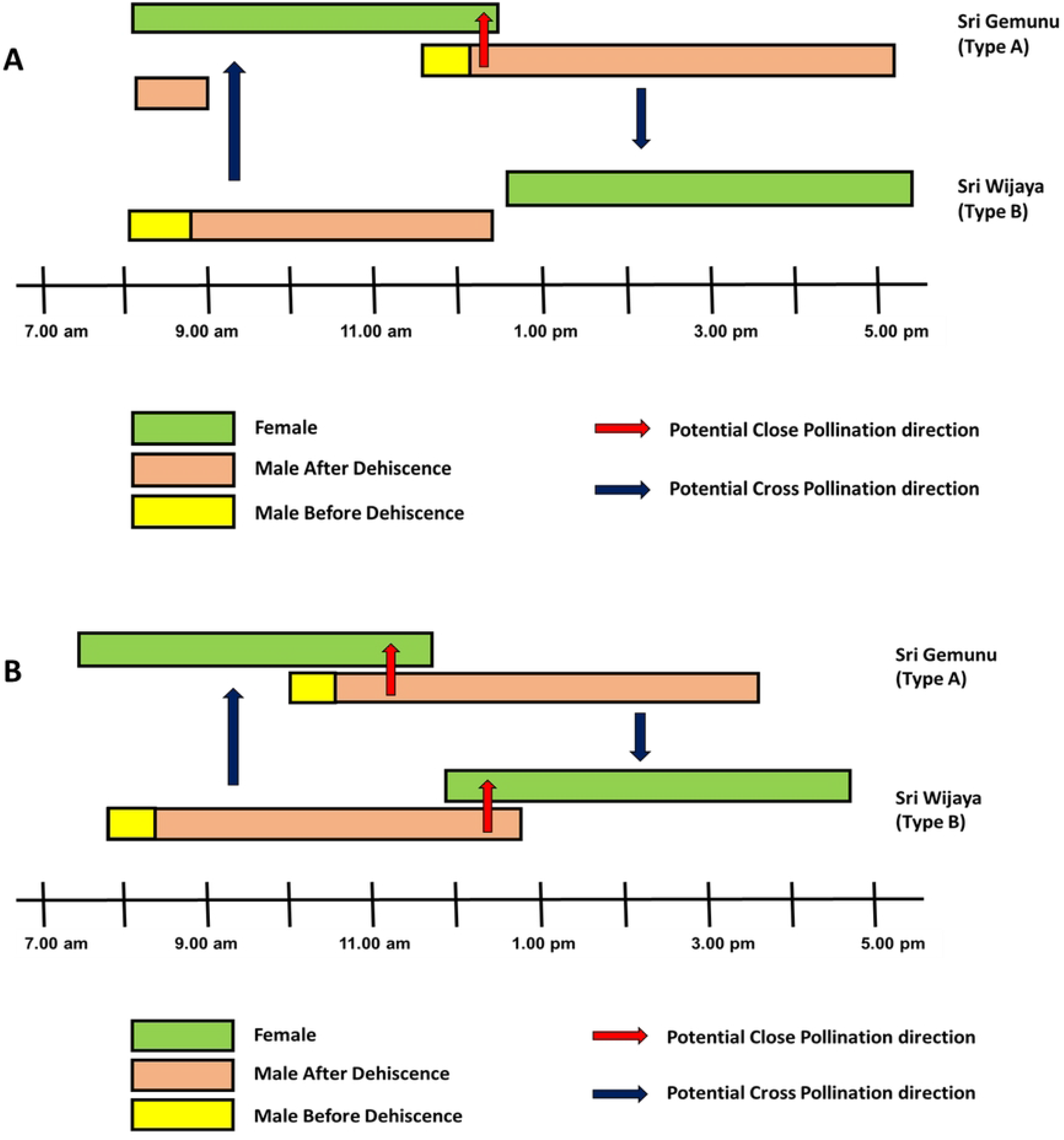
Floral behaviour of male and female flowers in the two verities under different envionemental conditions. (A) Flower opening sequence of *Cinnamomum verum (Sri Lankan)* under ideal temperature when all open flowers in a block are considered. The arrows indicate the potential pollen flow (B) Flower opening sequence in *Cinnamomum verum (Sri Lankan)* trees under cold weather conditions (temperature <25 °C) experienced in Delpitiya, Sri Lanka-when all open flowers in a block are considered. The arrows indicate the potential pollen flow.

During low temperature (maximum 25°C, minimum 22 °C) and high humidity (81.2%) (Supplementary Figs. 2,3) the flower opening and closing were delayed and extended (Fig 7 B). However, during cooler conditions, flower opening was delayed and extended. During cooler days, the opening of the *Sri Gemunu* male phase got delayed and then remained open till late evening (Fig 7 B). A similar trend was observed in the female stage of *Sri Wijaya* (Fig 7 B). Therefore, overlapping male and female stages on the same plant did not happen during the cooler weather.

Spearman’s rank correlation between the number of individuals in overlapping phases in both *Sri Gemunu* and *Sri Wijaya* is depicted in Table 2. There is a positive correlation between the flower overlapping with temperature in *Sri Gemunu* (r=0.484, p=<0.0001) and *Sri Wijaya* (r=0.477, p=<0.0001). It is negatively correlated to mean relative humidity in *Sri Gemunu* (r=-0.531, p=<0.0001) and *Sri Wijaya* (r=-0.535, p=<0.0001).

**Table 2:** Correlation of *Sri Gemunu* and *Sri Wijaya* floral overlapping with climatic factors.

|  | Environment Factor | Coefficient | t | DF | P value |
| --- | --- | --- | --- | --- | --- |
| *Sri Gemunu* | Mean relative humidity (%) | -0.531 | -13 | 430 | <0.0001* |
|  | Maximum daily temperatures (°C) | 0.484 | 11.47 | 430 | <0.0001* |
| *Sri Wijaya* | Mean relative humidity (%) | -0.535 | -9.03 | 214 | <0.0001* |
|  | Maximum daily temperatures (°C) | 0.477 | 7.95 | 214 | <0.0001* |

\*Significant at p<0.001

Table 3 shows the synchrony of phonological events of the two varieties *Sri Wijaya* and *Sri Gemunu*. The highest synchrony ratio indicates a greater coincidence of the phase among individuals. The overall inter-population synchrony ratio for flower overlapping between male and female flowers during the overlapping period is higher during the peak season. The overall synchrony ratio during peak season/mass flowering during January and February for *Sri Gemunu* and *Sri Wijaya* are 0.873± 0.040 and 0.854±0.026, respectively (Table 3).

**Table 3:** Synchrony indices for phenological events of *Sri Gemunu* and *Sri Wijaya* within Peak and off-peak season.

| Variety | Season | SI | temperature | humidity |
| --- | --- | --- | --- | --- |
| *Sri Gemunu* | Peak | 0.873± 0.040<sup>a</sup> | 30.83 | 59.3 |
|  | Off peak | 0.687±0.049<sup>b</sup> | 26.83 | 76.7 |
| *Sri Wijaya* | Peak | 0.854±0.026<sup>a</sup> | 30.83 | 59.3 |
|  | Off peak | 0.6239±0.031<sup>b</sup> | 26.83 | 76.7 |

### Receptivity of the stigma

The stigma supports pollen hydration, germination, and initial pollen tube growth, and the length of stigmatic receptivity plays an important role in the effective pollination period and subsequent fruit set [23,24]. The stigma receptivity continued up to the male stage since the female flower opening. The highest stigmatic receptivity was observed between 8.30 am to 10.30 am in *Sri Gemunu*, while for *Sri Wijaya* it was around 1.00 pm to 4.00 pm (Fig 8, Table 4). These results suggest that the *Cinnamomum* stigma maintains its capacity to support pollen adhesion and pollen germination during the length of the female phase. Thus, the pollen could be applied to the stigma along the female stage during hand-pollination although the stigma seems to provide higher adhesion during the middle part of this period.

**Fig. 8:**
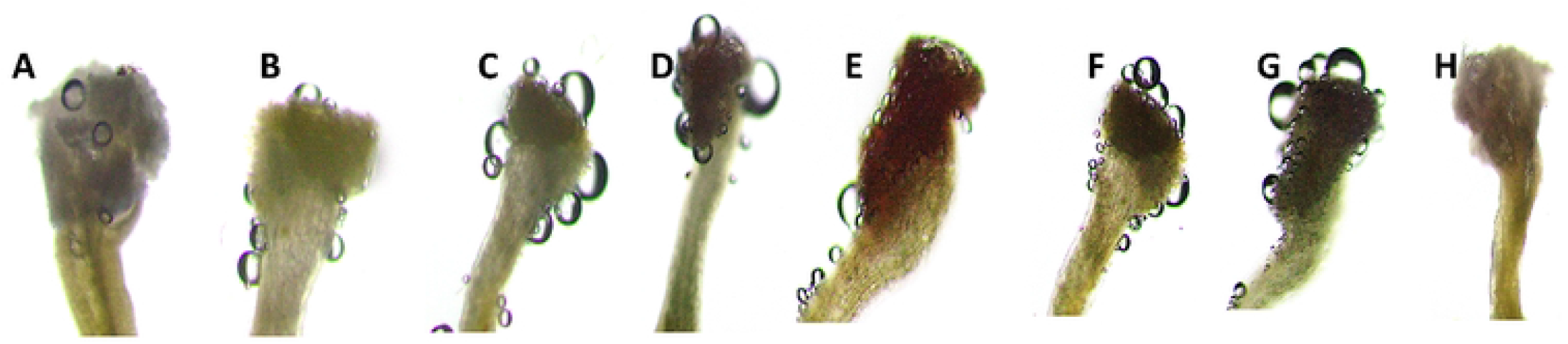
Evaluation of the stigmatic receptivity of *Cinnamomum verum* with hydrogen peroxide. (d-g); a) No reaction in pre-anthesis; b) Strong positive reaction (+) in anthesis; c) Very strong positive reaction (++) in after 2 hours; Very strong positive reaction (++) in after 2 hours d) Weak positive reaction (+) in pre-anthesis; e) Strong positive reaction in anthesis; f) Very strong positive reaction in anthesis; g) Very strong positive reaction in post-anthesis.

**Table 4:**
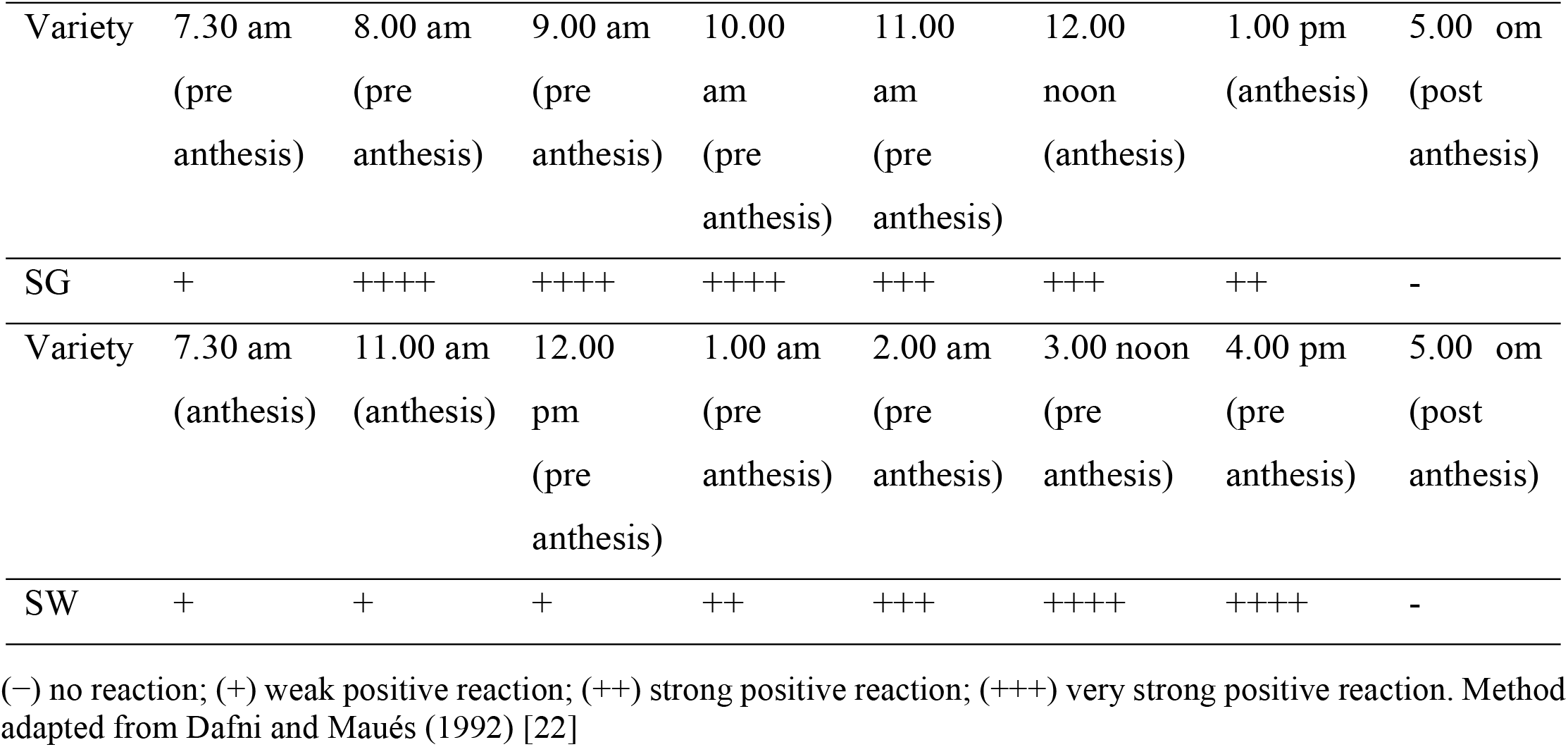
Stigma receptivity in *C. verum* of the two varieties *Sri Gemunu* (SG) and *Sri Wijaya* (SW) evaluated by hydrogen peroxide at different times related to anthesis.

| Variety | 7.30 am<br>(pre<br>anthesis) | 8.00 am<br>(pre<br>anthesis) | 9.00 am<br>(pre<br>anthesis) | 10.00<br>am<br>(pre<br>anthesis) | 11.00<br>am<br>(pre<br>anthesis) | 12.00<br>noon<br>(anthesis) | 1.00 pm<br>(anthesis) | 5.00 om<br>(post<br>anthesis) |
| --- | --- | --- | --- | --- | --- | --- | --- | --- |
| SG | + | ++++ | ++++ | ++++ | +++ | +++ | ++ | - |
| Variety | 7.30 am<br>(anthesis) | 11.00 am<br>(anthesis) | 12.00<br>pm<br>(pre<br>anthesis) | 1.00 am<br>(pre<br>anthesis) | 2.00 am<br>(pre<br>anthesis) | 3.00 noon<br>(pre<br>anthesis) | 4.00 pm<br>(pre<br>anthesis) | 5.00 om<br>(post<br>anthesis) |
| SW | + | + | + | ++ | +++ | ++++ | ++++ | - |

(–) no reaction; (+) weak positive reaction; (++) strong positive reaction; (+++) very strong positive reaction. Method adapted from Dafni and Maués (1992) [22]

### Pollen Morphology and Pollination Biology

The SEM images describe the morphology of the pollen grains of two Cinnamon varieties (Fig 9), and the quantitative and qualitative features are summarized (Table 5). The pollen grains of the investigated *C. verum* varieties are spheroidal in shape, apolar and inaperturate. As expected, both varieties have the same shape pollens (Fig 9 A, E). The exine ornamentation is spinulate with a spine having a conspicuous cushion base. This cushion base is prominent and ornamented with balls-like structures. According to Erdtman’s (Erdtman, 1986) pollen size classification, most of the investigated pollen grains were median with the polar axis (P) of corpus ranging from 22-24 μm. *Sri Gemunu* pollen is larger than that of *Sri Wijaya* in size. The spine length range from 0.8-to 0.9 μm and the spines of the *Sri Gemunu* is placed at a distance than the *Sri Wijaya (*Fig 9 B, F). The spinules of *Sri Wijaya* pollen are shorter than that of the *Sri Gemunu. T*he spinule tip of the *Sri Gemunu* pollen is pointed, conical shaped, and striated (Fig 9 C), while the *Sri Wijaya* are acute and blunt, conical linear, and striated (Fig 9 G). Moreover, the tectum is highly perforated in the *Sri Gemunu*, while it is less in the *Sri Wijaya* (Fig 9 D, H).

**Fig. 9:**
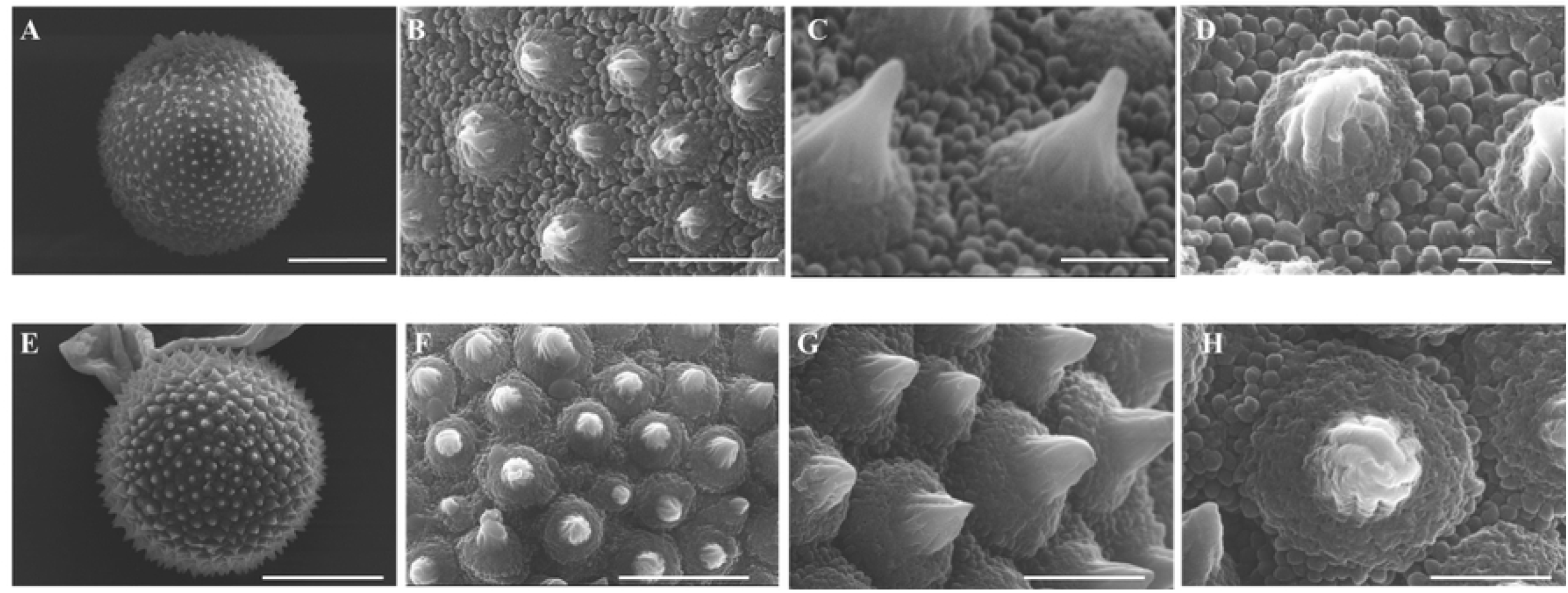
Scanning electron microscopy (SEM) images of pollen grains of *Cinnamomum verum (Sri Lankan);* Sri Gemunu and Sri Wijaya. (A,B,C,D) *Cinnamomum verum (Sri Lankan)* Sri Gemunu Type A (A) Polar view of pollen grains (B) detail of spines and exine structure /Exine ornamentation (C) Dorsal view of an enlarged spine, tectum is highly perforated (D) Ventral view of the enlarged spine, striated conical-shaped spine on a cushion base (E,F,G,H) *Cinnamomum verum (Sri Lankan)* Sri Gemunu Type A (E) Polar view of pollen grains (F) detail of spines and exine structure /Exine ornamentation, tectum is less perforated (G) Ventral view of a Striated Conical linear spine on a cushion base (H) Dorsoventral view of a spine

**Table 5:** Quantitative and qualitative pollen morphological characters of *Cinnamomum*.

| Quantitative |  |  |  |  |  |  |
| --- | --- | --- | --- | --- | --- | --- |
| | Polar length (P) $\mu\text{m}$ | Length of the spinule $\mu\text{m}$ | Length of the spinule with base | Interspinular Distance $\mu\text{m}$ | Diameter of the spinule base | |
| *Sri Gemunu* | 24.36±0.123 | 0.86±0.082 | 1.11±0.113 | 0.48±0.103 | 0.86±0.082 | |
| *Sri Wijaya* | 22.22±0.123 | 0.89±0.012 | 1.48±0.018 | 0.80±0.100 | 1.60±0.007 | |
| Qualitative |  |  |  |  |  |  |
|  | shape class of pollen | Tip of spines | Ornamentation of spines | Types of granules | Nature of granules | Nature of cushion base |
| *Sri Gemunu* | spheroidal | Acute | Striated | Monomorphic | Prominent | Prominent |
| *Sri Wijaya* | spheroidal | Acuminate-acute | Striated | Monomorphic | Prominent | Prominent |

### Floral visitor observations

According to the field observations, flowers of *C. verum* are assumed to be pollinated by insects including mainly Hymenoptera, Diptera Syrphidae, and Coleoptera. Namely, honeybees (*Apis mellifera, Apis cerana*), Hoverfly (*Dideopsis aegrota*), Wasps (*Delta dimidiatipenne*), bee (*Trigona iridipennis*), Butterfly (*Ypthima ceylonica*), and *Formicidae* sp (*Oecophylla smaragdina*), flies (Housefly *Lucilia cuprina*) and mosquitoes (Figure 10). They forage daily during day hours from 0800-1800h collecting pollen and nectar. When visiting flowers, bees grasped petals by using their feet to fix the body, generally staying on a single flower for about 10-30 seconds and then flying to another flower.

**Fig. 10:**
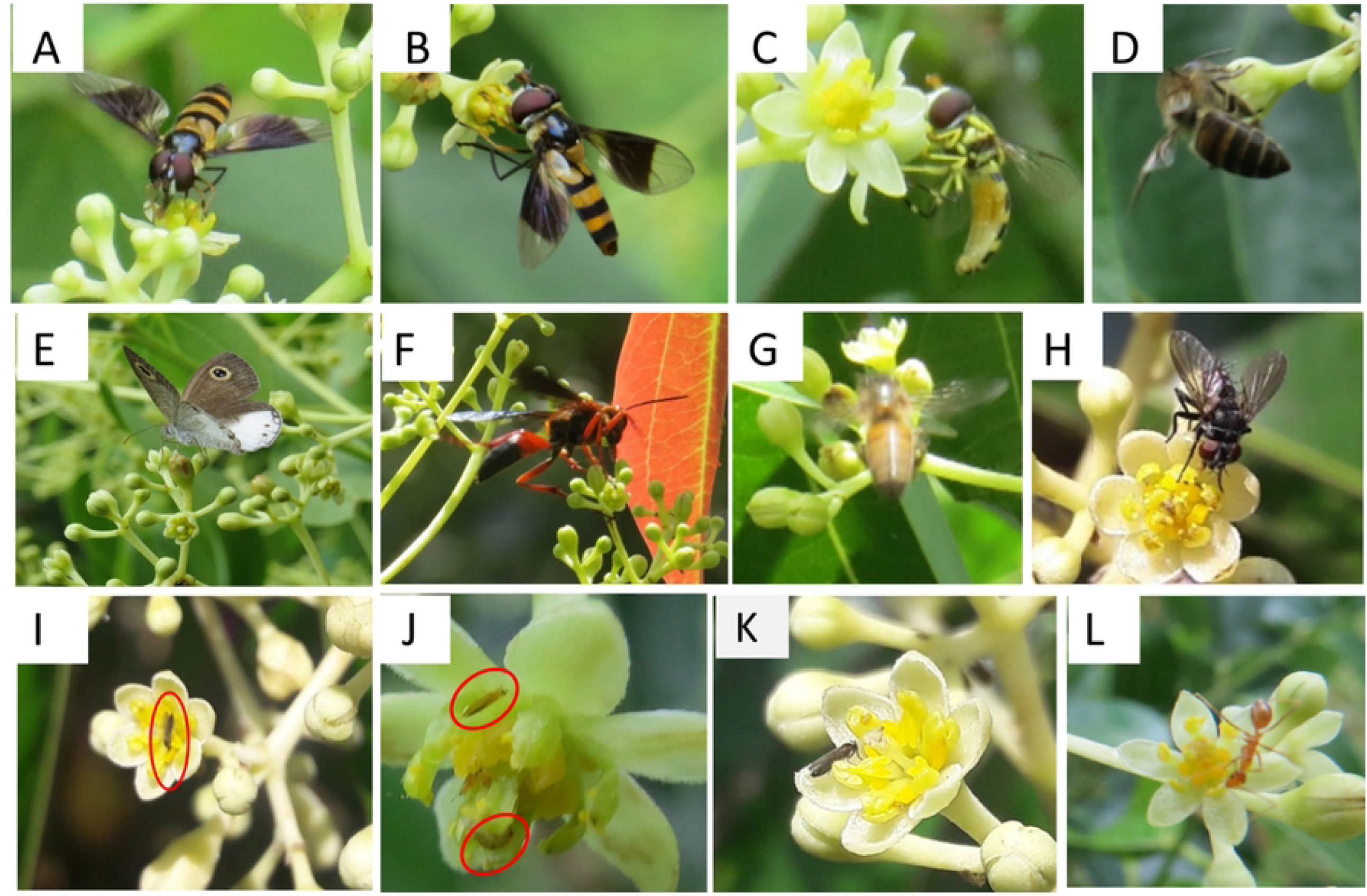
Pollinator’s foraging behaviors on Cinnamon flowers. (A-B)-Hoverfly/Syrphid fly or Flower fly-*Dideopsis aegrota* (C)-Hoverfly *Syrphus ribesii* (D)-Apidae Honeybee *Apis cerana* (E) butterfly *Ypthima ceylonica* (F) Wasp Delta dimidiatipenne (G) Apidae Honeybee Apis mellifera (H) House fly *Lucilia cuprina* (I) Small flies (J)-Thrips (K) Mosquiotes (L) Formicidae Oecophylla smaragd

The average duration of the visit made by hoverflies is 5-25 sec; *Wasps* with 1 minute between 10.00 and 18.00h, *Apis mellifera* visits for 8-25 sec between 09.00 to 13.00h. Butterfly visit for 2 to 5 sec during 09.00 to 12.00h and the duration of Ants visit is highest between 08.00 to 18:00h. The honeybees and the hoverflies are the dominant pollinators. Our observations suggest that excluding mosquitoes and thrips that may act as nectar thieves, other floral visitors were effective pollinators that could perform pollination while they were collecting pollen or sucking nectar.

### Reproductive output under controlled hand pollination

The percentage of fruit set in the three plants isolated are given in Table 6.

**Table 6:** Fruiting rate observed after controlled pollination systems.

| Plant | Status | Total number of flowers produced | Total number of fruits produced | Percentage of fruit set |
| --- | --- | --- | --- | --- |
| 01 | Insect pollinated/Plant house (higher temp/low humidity) | 46 | 2 | 4.32% |
| 02 | Hand pollinated | 148 | 12 | 8.01% |
| 03 | Isolated plant in an open area | 538 | 0 | 0% |

The fruiting rate was around 4% (Fig. 5.1 d) in both controlled insect-pollinated conditions and was around 8 % (Fig. 5.1 c) in hand-pollinated conditions. However, no fruiting was observed in isolated plant that was not controlled.

### Histological evidence for self-incompatibility

Initial microscopic observation suggested that in close-pollinated flowers, crossed by hand, the pollen was reaching the stigmatic surface of the pistil, belonging to the same tree, and started to germinate indicating their viability. For the enhanced resolution of pollen-stigma interaction up to the pre-fertilization stage, a further microscopic study was conducted with the pistils collected at different stages of the floral cycle in the female stage. Subsequent pollen tube growth showed a differential pattern between cross-pollinated and close-pollinated flowers to cover the stylar canal during the first 24 hours. The self-pollinated pistils developed cellulosic walls at and around nucleus cells surrounding the megaspore. This callose growth progressively increased with time (Fig 11 E, F, G), restricting reaching the ovule (Fig 11 H).

**Fig. 11:**
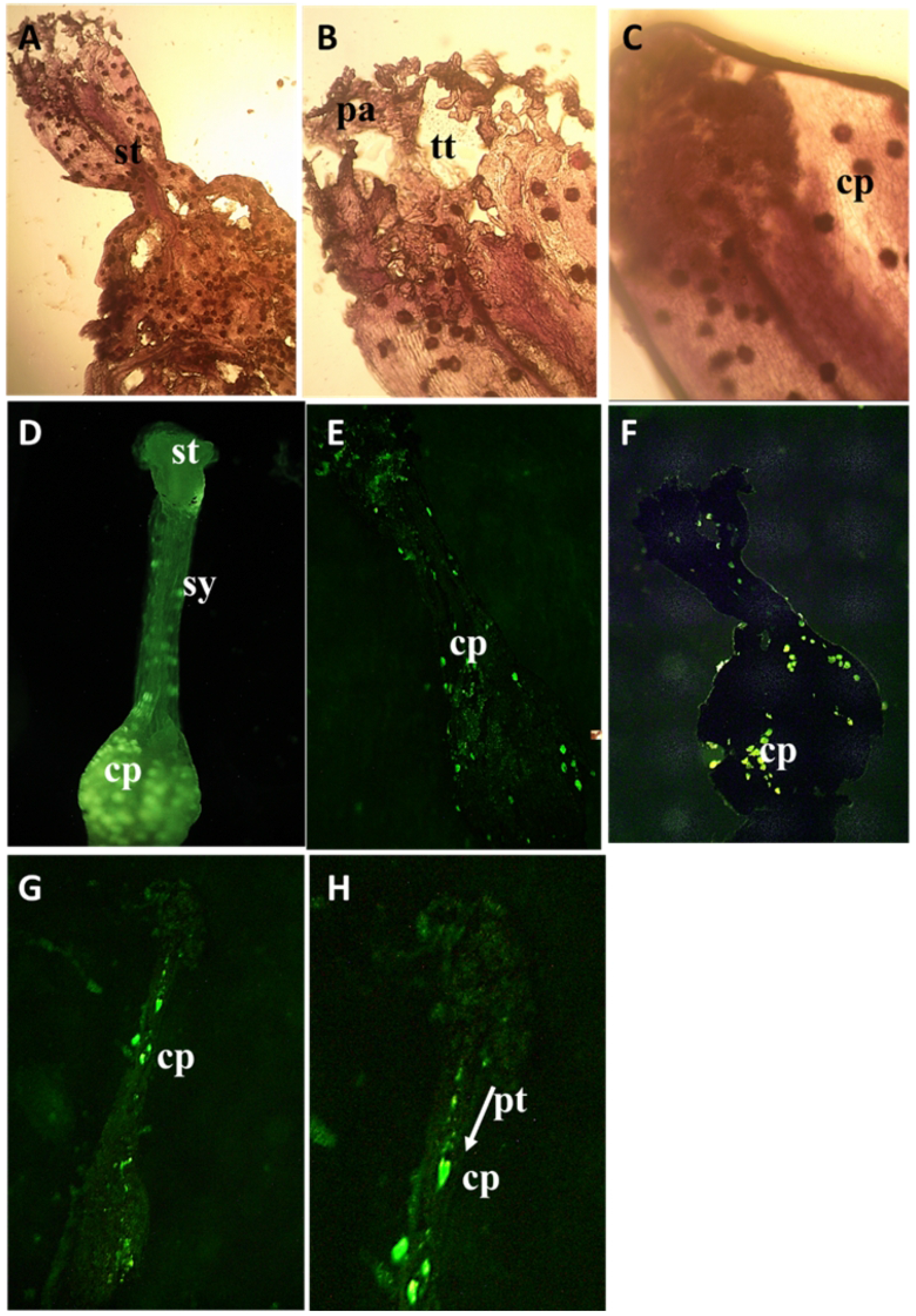
Pollen tube growth analysis in incompatible and compatible assisted crosses using fluorescence microscopy. Callose plugs (cp) have been formed so as to inhibit the pollen tube growth and fertilization (A,B,C) Light microscopic images showing the development of callose plugs (D) Close-pollinated stigma (sg) after 24 hours (E) Close-pollinated stigma showing callose plugs thickening on style and ovary area after 1 hour (F) Close-pollinated stigma showing callose plugs thickening on style (st) and ovary area after 5 hours.(G,H), aniline blue staining of self-pollinated pistil after 20 hours, showing pollen tubes within the transmitting tract and reaching the ovules. Callose plugs thickening has inhibited the growth of pollen tubes by deposition at the tube tip, arrow showing the development of pollen tube (pt)

During the pre-receptive period, the stigma was slightly enlarged (Fig 11 A), papillae did not expand completely, and some remained relatively wrinkled. During the receptive period from around 8.00 am to 11.00 am, the stigma starts secreting exudate from vesicles and secretions seen on the surface, and some papillae had periplasmic spaces where secretions may be present. During the post-receptive period, the stigma turned blackish, and the remnants of the exudate were found on some stigmata (Fig 11 B).

### Changes of Stigma Surface at Different Flowering

The SEM observations of the stigma soon after the opening showed a prominent slightly irregular triangular-shaped stigma (Fig 12 A), ending in a smooth elongated style. At the earliest stage of female opening, before the pollination, the trilobe-shaped stigma was bordered by rows of feathery appendages (Fig 12 B). These feathery-like structures are the raised unicellular papillae. The papillae were closely packed and formed ridges-like structures (Fig 12 C). After the stigma starts being receptive, the stigmatic papillae displayed round-shaped secretory vesicle-like structures fusing to the plasma membrane. These secretory vesicles cover the full surface for 30 minutes following compatible pollinations (Fig 12 D, E). At this time, a thin layer of stigmatic secretion from the secretary vesicle coated the papillary surface (Fig 12 F), and the stigma was visually glossy and milky white (Fig 2 A). During later stages and after pollination, the papillae became more distinct, and accumulated stigma exudate appeared at some interstitial regions at the base of the adjacent papillae. The pollinated stigmas naturally turned brownish-black (Fig 2 B). In some flowers, which appeared to be pollinated and germinated, the stigmatic surfaces collapsed forming a hollow-like thing in the middle (Fig 12 G).

**Fig. 12:**
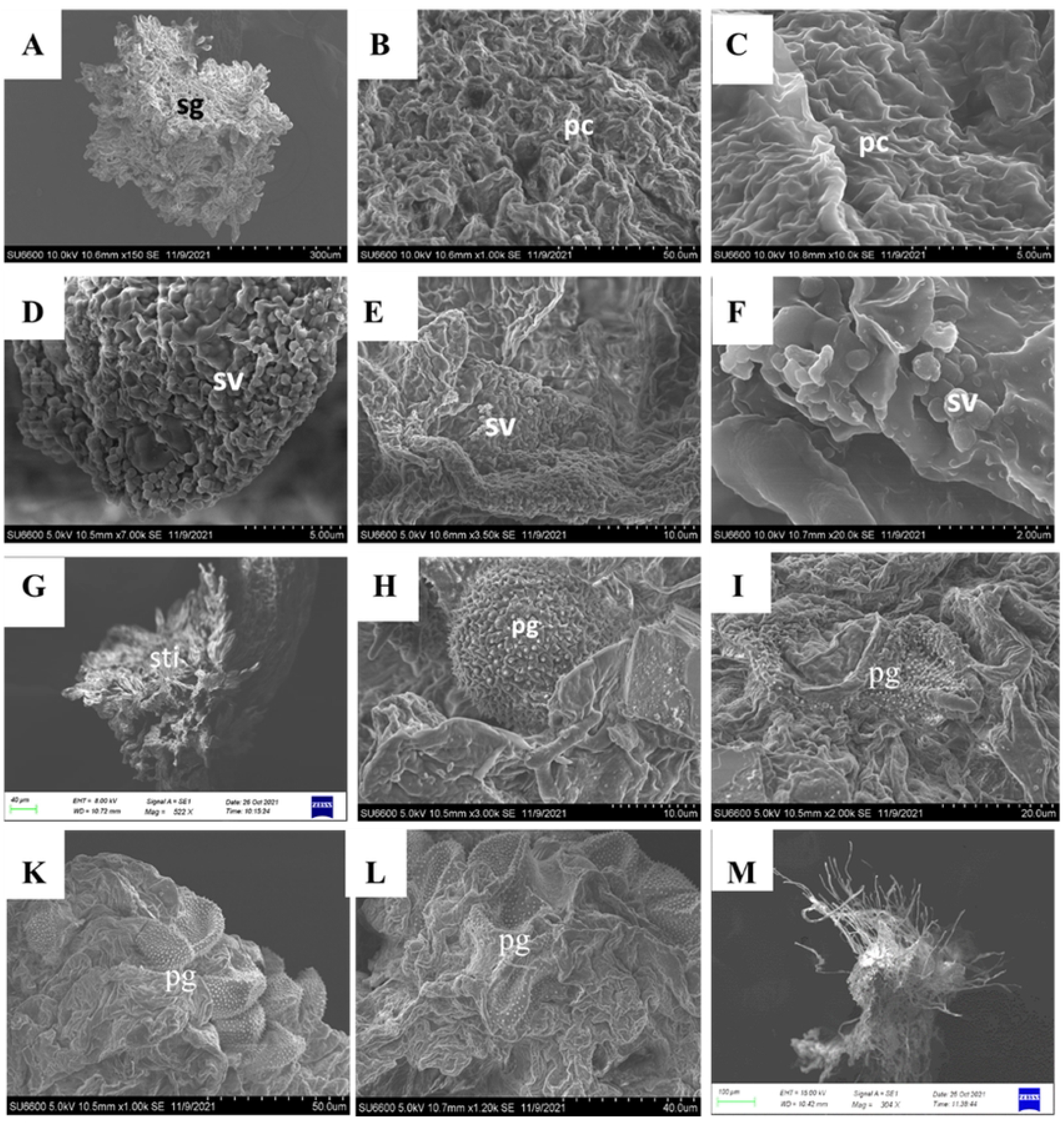
Stigma morphology variation in *Sri Gemunu* during the floral cycle during close pollination. (A) Dorsal ventral view of the fresh female stigma (sg) soon after opening (B) Stigmatic surface with elongated stigmatic papillary cells (pc) in ribs-like structures, providing higher surface area for pollen adhesion (C) Higher magnification of the papillary cell layer (D,E) Secretory vesicles (sv) appeared on the stigma after the stigma being receptive, the secretory vesicles are circular lobes-like structures, arranged all over the papillae cell layer (F) Higher magnification of secretory vesicles (G) The stigma has a central depression forming a short stigmatic cleft in the pollinated stigma, feather-like papillary cells are distinct (H) Hollen grains deposited on the female stigma, pollen merged on the stigmatic surface (I) Pollen grains deposited on the stigmatic surface shrank and shriveled after pollination (J,K) Magnified Functional male stigmatic surface with multiple pollen deposited, all those pollen grains merged into the stigmatic surface, the pollen grains are shrunk and shriveled. (L) Un-pollinated stigma in male flower, the papillae cells converted into fiber-like structures

During stigma receptivity, the surface area of the stigma increased considerably due to the physical appendage increase in the stigmatic lobes and enhanced the exertion of globular papillae on the surface. After opening the flowers for about 2 hours (pre-anthesis), the stigma surface was densely covered with round unicellular secretary vesicles. These numerous vesicles on the stigma surface provide a receptive surface for compatible pollen grains to adhere, hydrate, germinate and grow. Within minutes of the capture of grain by the stigma, the exine-held material flows out and becomes deposited on and between the stigmatic papillae. Pollen grains were attached at the end of the papillae surface, and some were deposited on the stigma surface (Fig 12 H).

After the anthesis, the stigma visually turned from milky white color into black. The pollinated stigma was visually abundant with pollen grains (Fig 12 K). Most of the pollen grains were shrunk, shrivelled, and drowned into the stigmatic surface through the papillary spaces (Fig 12 K, L). These shrunk pollen grains may be due to incompatibility. The pollen grains were not hydrated from the stigmatic surfaces and the papillae in the unpollinated stigma turned into fiber-like structures (Fig 12 M). The unpollinated stigma was separated from the style (Fig 12 I). There was no distinct difference between the stigmatic surfaces in *Sri Gemunu* and *Sri Wijaya* (Supplementary Fig 4), both varieties exert similar mechanisms in pollen deposition, hydrations, and germination.

## DISCUSSION

The synchronously protogynous flowering rhythm is accepted as a typical adaptation to cross-pollination. In *Cinnamomum*, pollen transfers from B-type cultivars to A-type cultivars during the morning and vice versa in the afternoon. Our observations and data suggest classical protogynous dichogamy behaviour with spatial and temporal separation of male and female functions in *C verum*. The biological reason for special separation is to promote cross-pollination [14,25]. Every flower stage exhibits different structures and nectar, which are efficient for pollinator attraction [26]. The flowers are small and mostly bisexual and typically consist of seven trimerous whorls of floral organs; two perianth whorls, four androecial whorls, and one single carpel. Typically, the two perianth whorls are similar. Exceptionally abnormalities have been recorded by Azad in 2018 [27] like in two whorls and 7 in two whorls. The gynoecium in the functional female stage is considered as a single carpel and superior in most of the Lauracea [28,29]. The stamens of the third whorl bear a pair of appendages at the base. In the functional male stage, pollen sacs open introrsely in the two outer stamen whorls, while the third inner whorl opens extorsively. Anthers possess wither four pollen sacs in superimposed orientations similar to other members in Lauraceae [6,30]. However, in male flowers of *Cinnamomum*, all stamens bear paired staminal appendages.

The floral arrangements in cinnamon have generally been proposed to be adaptive mechanisms to limit self-fertilization and eliminate physical interference between pollen dispersal and receiving within flowers. The natural variation in inflorescence architecture also influences pollination behaviour and reproductive success, suggesting that inflorescence architecture can evolve under pollinator-mediated natural selection in plant populations [31–33].

In addition to the morphoanatomy mechanisms exerted for cross-pollination, rewards are readily available to floral visitors such as bees, wasps, ants, flies, beetles, bugs, and butterflies. However, effective cinnamon pollinators have to visit both female and male stage flowers and come in contact with the dehisced anthers and the receptive stigma at the hairy ‘pollen collection zones’. The flying insects of small to medium size (3 to 8 mm in length) are especially suitable for efficiently collecting cinnamon nectar [9]. The main cinnamon pollinators are the Hymenoptera species, several small to medium size stingless bees (Meliponinae), and honeybees. *Cinnamomum sulphuratum* and avocado, having similar pollination behaviour, consider honeybees the potential pollinator [34]. Second-order pollinators are numerous species of wasps, butterflies and flies and probably beetles, too [34]. These pollinators promote cross-pollination since they have a high chance of moving among neighbouring flowers and flying between inflorescences. However, it is considered that their pollination efficiency is not high because of a low rate of flower visitation [35,36]. Flies are one of the most abundant insects detected in the pollinator profile of cinnamon. In avocado, which has a similar type of floral biology, it has been identified that the thrips (*Frankliniella* and *Thrips palmi*) play a key role in pollination [37–39]. They are common visitors of cinnamon flowers during bloom [40]. Almost every flower has at least one trip, which freely roams the stamens and pistils. Thrips are a common insect pest in cinnamon that results in a significant yield loss in mature cinnamon plantations while it causes severe growth retardation in nurseries and younger plants [41]. The effectiveness of these insects for pollinating cinnamon flowers could be studied further in a closed field pollination environment. It is recommended to have a mixed plantation system of the two types of varieties to increase the cross-pollination probability. It will also reduce the potential negative impact of a lack of synchrony between the two varieties caused by environmental factors.

The physiological and structural originations of the floral organs also maximize cross-pollination. Assessing the morpho-anatomy and stigma receptivity of cinnamon is vital in understanding the biological process. According to our observations, *Cinnamomum* stigma has papillae on its surface, and it develops secretory vesicles during the stigma receptivity period. No exudate was observed on the stigma surface at the flower opening, suggesting that *Cinnamomum* has a dry stigma, previously observed in plants having saprophytic self-incompatibility systems [42,43].

Pollen receptivity is dependent on the nature of the stigmatic surface [44]. Moreover, the amount of pollen grains received by the stigma is influenced by its surface area. Accordingly, *C. verum* has a unique stigma surface in the female stage compared with the other species in the family. The receptive stigmatic dry surface is concentrated in distinct ridges with unicellular papillae, increasing pollination efficiency. This arrangement could prevent freshly deposited pollen grains from falling off the stigma and possibly a precaution for the pollen tube growth from close-pollination. The numerous stigma feathers-like structures in cinnamon create a relatively larger area that increases the probability of pollen grain deposition [45,46]. The same has been recorded in avocados [28,47]. The secretory vesicles distinct after the stigma starts being receptive would be a critical factor affecting the pollen receptivity in stigma, making it wet to bind the pollen. Close-pollination barriers on the stigma surface result in the arrest of pollen germination or pollen tube entry into the stigma [48]. The causative factors for the failure of pollen germination can be due to the lack of effective adhesion, lack of full hydration and pollen germination factors on the stigma [42,49–51]. Pollen adhesion largely depends on the nature and extent of the stigma surface, specifically for dry-type stigma [43,48,52,53].

Normally, compatible pollens get hydrated due to the transfer of water from the stigma to the pollen through an osmotic gradient. However, according to our SEM images, pollen shrinkage and shrivelling were prominent during close pollination, which may be due to insufficient or uncontrolled hydration. Even after effective adhesion, pollen grains require suitable conditions on the stigma for germination. Focusing on the stigma itself after anthesis, our observations suggest that pollen grains are drawn into the stigmatic surfaces through the papillary spaces. Pollen tube growth on *Cinnamomum* is not surficial, or even immediate subcuticular, as in stigmatic papilla of *Persea americana* [47,54] and other angiosperms with dry stigma including Brassica [48,55] and Nymphaea [56]. It occurs deeper within the stigmatic cells. The stigmatic papillae of *Cinnamomum* are morphologically different from those of avocado [57,58], in terms of stigmatic vesicles. Those were visually different from other dry stigmas. Thus, *C. verum* could be a useful model for comparative studies in pollen-pistil interactions in the Lauraceae family.

Moreover, the formation of callose plugs at and around the style and ovary is clear evidence of its self-incompatibility. These callose plugs form in close-pollinated pistils and may be the result of such a self-recognition mechanism [59]. The development of such mechanical barriers in growing pollen tubes in pistils via long-distance signalling mechanisms has been extensively discussed in other families like Brassicaea [42,47,60,61]. It was evident that the size of the callose plugs increased over time, which could result in the degeneration of ovules in the close-pollinated pistil. Callose plugs observed at and around the nucleus cell/ovary may prevent nutrients from reaching the developing megaspore mother cell/embryo [51,60].

In contrast to all these adaptations; cross-pollination and self-incompatibility systems, some deviations were observed, from close pollination. We achieved ∼8% fruit set in *Cinnamomum* plants in a single closed environment by hand pollination, while insect pollination resulted in ∼4% productivity. This overlapping is more extensive at the high temperature and low humidity in peak and off-peak seasons. Increasing the overlapping time between flowers in both male and female stages of the same plant or plants from the same variety could be a strategy to ensure close pollination. Moreover, controlling temperature and humidity extended the stigma receptivity and the overlapping period. Extended stigma receptivity assures pollination, fertilization, and reproductive success in plants [24,43,62,63].

Ants could help in close pollination, and it has been identified in other *Cinnamomum* species as well [34]. Supporting the field observations, the SEM analysis showed that there is also a possibility of pollen grain germination in a relatively low percentage during close pollination. Hydrated pollens deposited on the stigmatic papillae were observed. This is supported by a recent field study [64]. Of the Sri Wijaya offspring resulting from a single open-pollinated event, 20% of individuals shared identical ISSR fingerprints with the mother plant.

Accordingly, *C. verum* has a complex reproductive mechanism, which may be a reason for the huge morphological, biochemical and genetic diversity observed. More focused breeding strategies are needed for crop improvement to cope with increasing demand and biotic and abiotic challenges including climate change. For example, close pollination within the tree is recommended to produce an elite seedling progeny from identified with superior mother plants or accessions. The grower will be able to enhance the seed setting under a controlled pollination environment with low temperature and high humidity.

## Acknowledgements

The authors acknowledge the Department of Export Agriculture, Peradeniya, Staff of the Mid Country Sub Research Station (MCRS), *Delpitiya*, for providing permission for carrying out the field experiments and flower collection. We would also thank the Analytical Department of the Sri Lanka Institute of Nano-Technology, Pitipana, and Homagama for capturing the SEM images. The funders had no role in study design, data collection and analysis, decision to publish, or preparation of the manuscript.The authors would like to thank the staff of the Agricultural Biotechnology Centre, Faculty of Agriculture, University of Peradeniya, for the support extended throughout the period.

## Author Contributions

**B.M.H** data curation, formal analysis, methodology, software, validation, visualization Writing – original draft and editing, **D.K.N.G.P**. Supervision, conceptualization, validation, visualization and writing-review, **P.C.G.B**. Supervision, conceptualization, funding acquisition, investigation, resources, visualization and writing –original draft and review

## Data Availability Statement

All the data available without fully restriction. Supplememtary data submitted.

## Supplementary Information

**Supplementary Fig. 1: SEM images of dehisced stamens in male flower** (A) Lateral view f Stamen Fourth whorl (S4) (B) First-whorl stamen at anthesis (with open stoma flaps); mature pollen grains at the opened pollen sac and the filament with fewer trichomes (C) Valvular dehiscence from the first whole stamen, Mature pollen grains distinct in opened pollen sac

**Supplementary Fig. 2:** Progression of overlapping percentage in *Sri Gemunu* of both female and male flowers during the overlapping period in peak and off peak season, with temperature, and humidity a-f G1,G2,G3,G4,G5,G6

**Supplementary Fig. 3:** Progression of overlapping percentage in *Sri Wijaya* of both female and male flowers during the overlapping period in peak and off peak season, with temperature, and humidity a-f W1,W2,W3,W4,W5,W6

**Supplementary Fig. 4: Stigma morphology variation in *Sri Wijaya* during the floral cycle**

**(A)** Lateral view of the fresh female stigma soon after opening (B) Dorsal ventral view of the fresh female stigma (sg) soon after opening (C) Stigmatic surface with elongated stigmatic papillary cells (pc) in ribs-like structures, providing higher surface area for pollen adhesion (D,E) Secretory vesicles (sv) appeared on the stigma after the stigma is receptive, the secretory vesicles are circular lobes-like structures, arranged all over the papillae cell layer (F) Pollen grains deposited on the female stigma, pollen merged on the stigmatic surface in the female flower (G) Pollen grains submerged on the stigmatic surface for the pollen tube growth, after pollination (H) Magnified Functional male stigmatic surface with multiple pollen deposited, all those pollen grains merged into the stigmatic surface, the pollen grains are shrunk and shrivelled. (I) Pollen grain submerged on the stigmatic surface for pollen growth detected in male stigma (J) Filaments observed in the style in the male flower stigma (K) Central groove observed in the style after pollination (L) The stigma has a central depression forming a short stigmatic cleft in the pollinated stigma, feather-like papillary cells are distinct

